# Arrestin-3 agonism at D3 dopamine receptors defines a subclass of second generation antipsychotics that promotes drug tolerance

**DOI:** 10.1101/2022.08.17.504324

**Authors:** Selin Schamiloglu, Elinor Lewis, Anne C. Hergarden, Kevin J. Bender, Jennifer L. Whistler

## Abstract

Second generation antipsychotics (SGAs) are front-line treatments for serious mental illness. Often, individual patients benefit only from some SGAs and not others. The mechanisms underlying this unpredictability in treatment efficacy remain unclear. All SGAs bind the D3 dopamine receptor (D3R) and are traditionally considered antagonists for dopamine receptor signaling. Here, we report that some clinically important SGAs function as arrestin-3 agonists at D3R, resulting in modulation of calcium channels localized to the site of action potential initiation in prefrontal cortex pyramidal neurons. We further show that chronic treatment with an arrestin-3 agonist-SGA, but not an antagonist-SGA, abolishes D3R function through post-endocytic receptor degradation by G-protein coupled receptor-associated sorting protein-1 (GASP1). These results implicate D3R-arrestin-3 signaling as a source of SGA variability, highlighting the importance of including arrestin-3 signaling in characterizations of drug action. Furthermore, they suggest that post-endocytic receptor trafficking that occurs during chronic SGA treatment may contribute to treatment efficacy.

## INTRODUCTION

Second generation antipsychotics (SGAs) are important tools in the management of serious mental illness (SMI), which includes bipolar disorder, major depressive disorder, schizophrenia, and schizoaffective disorder (Chouinard et al., 2017; Horacek et al., 2006). Each of these medications has unique dosing, pharmacokinetics, effect/side effect profiles, and cost, and clinicians must often prescribe several drugs before finding a treatment regimen that suits individual patients (Nivoli et al., 2011; Stahl and Grady, 2004). This approach primarily reflects an incomplete understanding of the molecular mechanisms underlying variable treatment efficacy in patients, even though these drugs have been in use for decades (Green and Braff, 2001). All SGAs are antagonists or partial agonists for G-protein signaling at the D2 dopamine receptor (D2R), and their clinical efficacy is thought to rely on their ability to block dopamine signaling at D2R (Howes and Kapur, 2009; Seeman, 2013). However, many SGAs also have high affinity for the D3 dopamine receptor (D3R) (Sokoloff and Le Foll, 2017), and the positron emission tomography (PET) experiments in human patients that form the foundation of the D2R antagonism hypothesis of SGA efficacy did not distinguish between D2R and D3R occupancy (Howes et al., 2009). Though D3R are more sparsely expressed than D2R, they are highly enriched in brain regions known to be altered in SMI, including striatum (Gurevich, 2022), Islands of Calleja (Zhang et al., 2021), and prefrontal cortex (PFC) (Smucny et al., 2022), where they define a unique population of layer V pyramidal cells (Clarkson et al., 2017). Furthermore, given these expression patterns in limbic systems, it is important to determine the effects of SGAs on D3R function (Sokoloff et al., 2006).

Dopamine receptors, like most G-protein coupled receptors (GPCRs), not only engage G-proteins upon activation, but also recruit other effectors, including arrestin-3 (β-arrestin 2) (Beaulieu and Gainetdinov, 2011; Urs et al., 2017). Arrestin-3 recruitment both “arrests” the canonical G-protein signal and also scaffolds non-canonical kinase activity, including Extracellular Signal Regulated Kinase 1 and 2 (ERK) (Beom et al., 2004). We recently demonstrated that ERK signaling via arrestin-3 is required for the ability of D3R to modulate calcium channel Ca_V_3.2 at the axon initial segment (AIS) after agonist activation (Yang et al., 2016). There has been no previous assessment of SGA ability to recruit arrestin-3 to D3R, despite the importance of D3R in SGA activity. Here, we report that some clinically important SGAs are “arrestin-biased” agonists at D3R, promoting recruitment of arrestin-3 and ERK activation in the absence of G-protein activation. We further demonstrate that only the arrestin-biased SGAs modulate Ca_V_3.2 activity in layer V neurons of the PFC. These findings provide a novel means by which SGAs can be divided into two functional classes based on their acute effects.

Importantly, the full therapeutic effect of SGAs often takes weeks to months of treatment. This suggests that mechanisms other than acute signal modulation via target receptors contribute to drug effect. In addition to its acute signaling roles, arrestin-3 also facilitates endocytosis (internalization) of GPCRs through interaction with protein components of the clathrin-coated pit (Shenoy and Lefkowitz, 2003; Zhang et al., 2016). Following endocytosis, both D2R and D3R are targeted for degradation in the lysosome via interaction with the GPCR-associated sorting protein-1 (GASP1) (Bartlett et al., 2005; Thompson and Whistler, 2011). Therefore, agonists that drive endocytosis decrease receptor surface levels over time, while conversely, antagonists that block both G-protein and arrestin-3 recruitment prevent dopamine-mediated endocytosis and thereby maintain receptor levels over time (Madhavan et al., 2013; Thompson et al., 2010). We hypothesized that chronic treatment with an arrestin-biased SGA would decrease the number of functional D3R and would do so with a time course more aligned with observed SGA treatment efficacy. We demonstrate here that chronic treatment with an arrestin-biased SGA eliminates the acute effects of D3R on AIS Ca_V_3.2 channel function in wild type (WT) mice but not in mice with conditional deletion of GASP1. We further demonstrate that mice treated chronically with an arrestin-3 biased SGA develop tolerance to the locomotor inhibitory effects of the drug and that this tolerance is abolished in mice lacking GASP1 in D3+ neurons. These findings provide a new mechanistic framework for understanding the therapeutic actions of SGA drugs.

## RESULTS

### Quinpirole modulates AIS calcium through D3R and arrestin-3 in PFC

The axon initial segment (AIS) is enriched with a number of ion channel classes, including Ca_V_3.2 calcium channels that are the target of neuromodulation via D3R (Clarkson et al., 2017; Dumenieu et al., 2018; Fukaya et al., 2018; Hu and Bean, 2018; Lipkin et al., 2021; Martinello et al., 2015). D3R-dependent modulation hyperpolarizes steady-state voltage-dependent inactivation of Ca_V_3.2 in a PKC- and ERK-dependent manner, leading to a reduction in high-frequency bursts of action potentials (APs) that depends in part on Ca_V_3.2 activity (Bender et al., 2010; Clarkson et al., 2017; Yang et al., 2016). In cartwheel cells of the dorsal cochlear nucleus (DCN), we showed that this effect requires both D3R and non-canonical arrestin-3 signaling (**Fig. 1A**) (Bender et al., 2010; Yang et al., 2016). We also reported previously that quinpirole, a D2/3R agonist, modulates Ca_V_3.2 at the AIS specifically in D3R-expressing (D3+) pyramidal cells in PFC (Clarkson et al., 2017). To examine whether this D3R-Ca_V_3.2 modulation at the AIS in PFC was also mediated by arrestin-3, as it was in DCN, we performed whole-cell current-clamp recordings from D3+ PFC pyramidal neurons in wild type (WT), D3R knock-out (D3 KO), and arrestin-3 knock-out (arrestin-3 KO) mice. D3+ neurons were identified either by fluorescent labeling of D3+ neurons in D3-Cre::Ai14 mice or, in tissue without fluorescence, by targeting neurons with specific intrinsic electrophysiological properties that allow for unambiguous identification of D3+ pyramidal cells (Clarkson et al., 2017). Using this approach (**Fig. 1B**), we replicated our previous findings that AIS calcium was reduced by 30.4 ± 2.9% after 20 minutes of quinpirole treatment (**Fig. 1C-D**) compared to time-locked vehicle controls (**Fig. 1D**). There was no change in AIS calcium in slices from D3 KO mice or arrestin-3 KO mice treated with quinpirole, and both were significantly different than WT slices treated with quinpirole (**Fig. 1D**). Therefore, in PFC, quinpirole-mediated modulation of AIS calcium in D3+ neurons also requires arrestin-3 (**Fig. 1A**), indicating that quinpirole is an agonist for arrestin-3 signaling at D3R.

**Figure 1:**
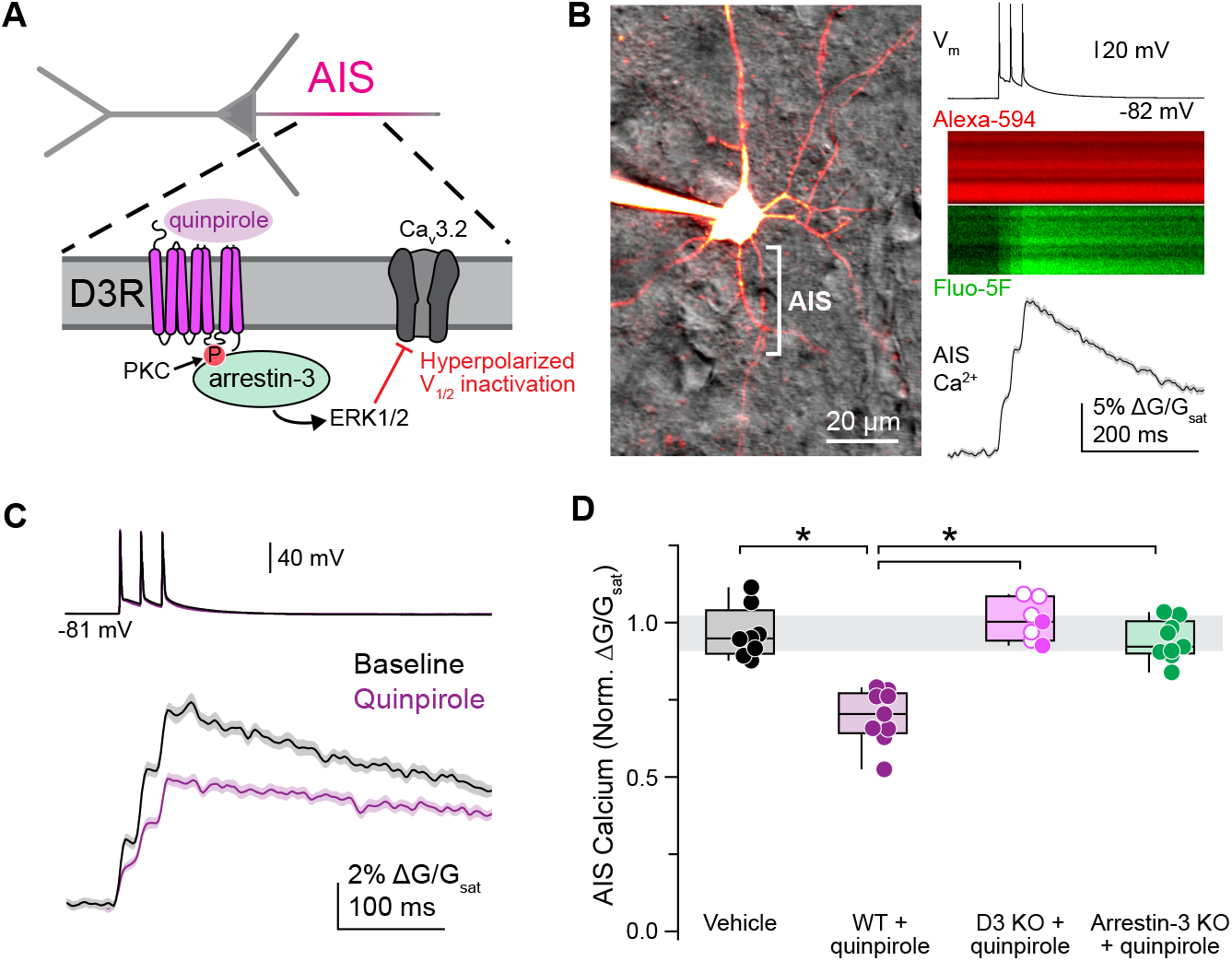
Quinpirole modulates AIS calcium through D3R and arrestin-3 in PFC. **A**. Schematic of quinpirole modulation of calcium at the AIS of a D3+ pyramidal cell in PFC. **B**. Left, 2PLSM z-stack of a D3+ pyramidal neuron visualized with Alexa Fluor 594. AIS denoted with bracket. Right, example linescan of AIS calcium averaged over 40 trials. APs were evoked with somatic current injection (3 at 50 Hz, 2 nA, 2 ms per stimulus). Linescan data displayed as mean ± SEM. **C**. Representative effect of quinpirole on AIS calcium in a D3+ neuron, averaged over 20 trials per condition. Baseline, black; quinpirole, purple. **D**. Peak AIS calcium transient amplitude normalized to baseline for vehicle and quinpirole in WT, D3 KO, and arrestin-3 KO mice. Vehicle controls include all 3 genotypes. For D3 KO, open circles denote previously published data (Clarkson, et al. 2017). Gray bar represents 95% confidence interval of control data. Vehicle: median normalized peak ΔG/G_sat_ = 94.8% of baseline, interquartile range (IQR) 89.9-104.0%, n = 8 cells from 6 mice; WT+quinpirole: 70.4% of baseline, IQR 64.2-77.2%, n = 9 cells from 7 mice, p <0.001; D3 KO+quinpirole: 100.3% of baseline, IQR 94.2-108.5%, n = 7 cells from 4 mice, p = 0.001; arrestin-3 KO+quinpirole: 92.2% of baseline, IQR 90.0-100.5%, n = 9 cells from 5 mice, p < 0.001. Kruskal-Wallis test with Mann-Whitney U test post hoc (Holm-Šídák correction).

### Arrestin-3 recruitment to D3R is both ligand- and PKC-dependent

Previous work has demonstrated that maximal endocytosis of D3R requires both receptor activation by agonist ligand and phosphorylation of the receptor by PKC (Cho et al., 2007; Thompson and Whistler, 2011). We hypothesized that arrestin-3 recruitment would also depend on both PKC activation and agonist binding. Consistent with this hypothesis, D3R-dependent modulation of AIS Ca_V_3.2 is both ligand and PKC-dependent, (Bender et al., 2010; Yang et al., 2016). We therefore examined whether PKC activity, concomitant with D3R agonist, is required for arrestin-3 recruitment to membrane D3R. To test this, we visualized changes in the distribution of cytosolic GFP-tagged arrestin-3 relative to FLAG-tagged D3R on the plasma membrane in HEK293 cells under a series of treatment conditions (**Fig. 2**). In untreated cells, and in cells treated with only the D3R agonist quinpirole or only the PKC activator phorbol-12-myristate-13-acetate (PMA), arrestin-3 remained diffusely distributed and did not colocalize with D3R at the plasma membrane (**Fig. 2B-C**). By contrast, arrestin-3 redistributed in cells treated with both quinpirole and PMA, moving to close proximity to the surface D3R (**Fig. 2B-C**). In quinpirole or PMA alone, peak FLAG and GFP signals remained well-separated (**Fig. 2B-C**), indicating that arrestin-3 was not recruited to membrane D3R, whereas co-application of quinpirole and PMA promoted redistribution of arrestin-3-GFP to the membrane. Under these conditions, FLAG and GFP fluorescence largely overlapped (**Fig. 2B-C**). Note that this timepoint preceded any significant receptor endocytosis. These results demonstrate that, in HEK293 cells, quinpirole is an agonist for arrestin-3 recruitment only with coincident PKC activation.

**Figure 2:**
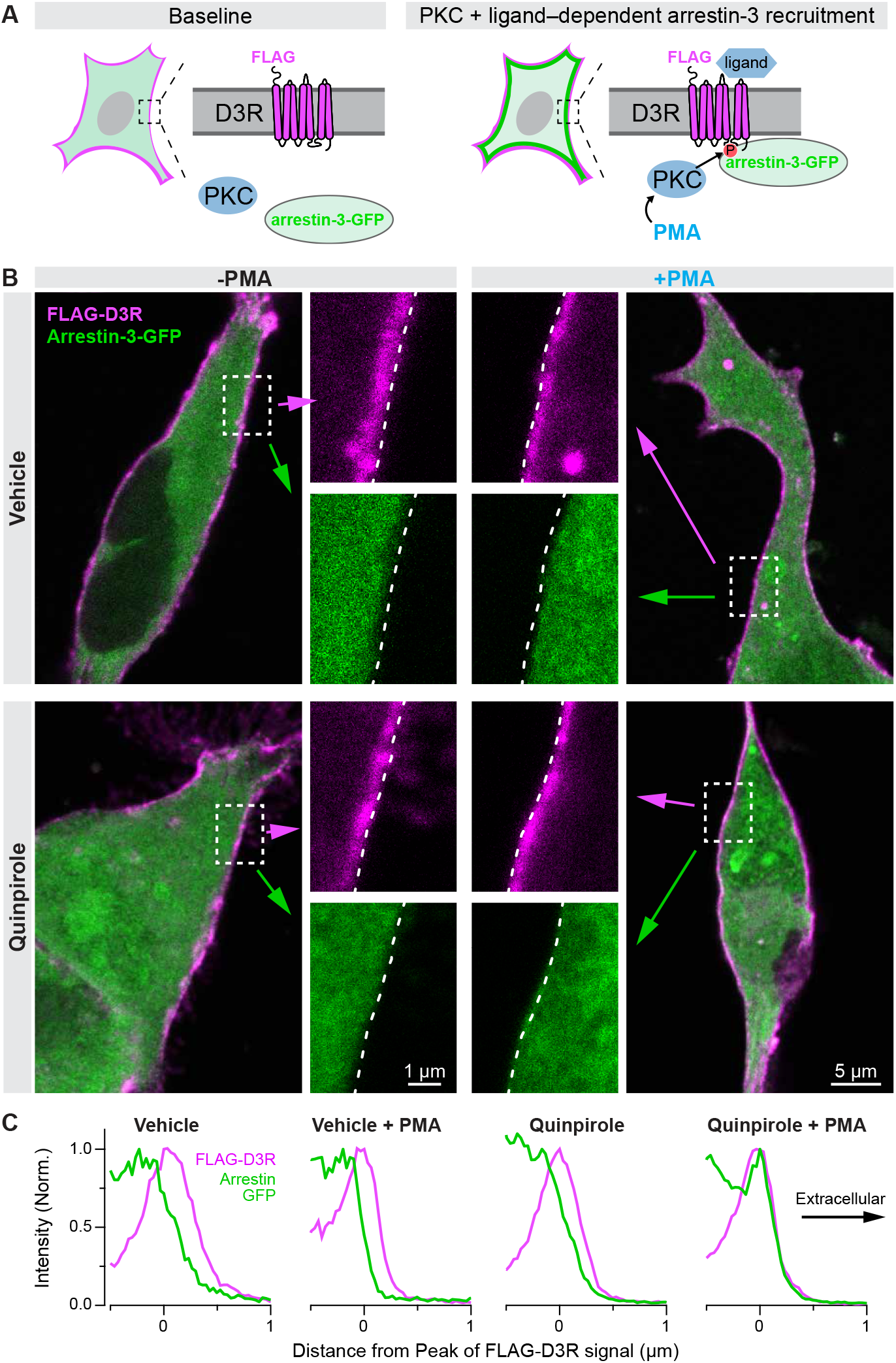
Arrestin-3 recruitment to D3R is both ligand- and PKC-dependent. **A**. Schematic showing that D3R ligand and PKC activation are both necessary to recruit arrestin-3 to D3R in HEK293 cells. **B**. Sample images of HEK293 cells expressing FLAG-D3R and arrestin-3-GFP. Cells were treated with quinpirole (bottom row) and/or PMA (right column). Dotted lines in insets denote edge of cell membrane. **C**. Quantification of images in B showing locations of peak FLAG-D3R and arrestin-3-GFP signal. FLAG-D3R in pink, arrestin-3-GFP in green.

### Some SGAs recruit arrestin-3 to D3R

There is mounting evidence that different G-protein coupled receptor (GPCR) drugs can stabilize distinct receptor conformations that preferentially engage either G-protein or arrestin-3 (Wootten et al., 2018). Based on this, we hypothesized that some SGAs could promote arrestin-3 recruitment to D3R even in the absence of G-protein activation. To examine this hypothesis, we performed arrestin-3-GFP redistribution experiments in D3R expressing cells (**Fig. 2**), but now with four commonly prescribed SGAs instead of the G-protein agonist quinpirole: roxindole, aripiprazole, quetiapine and clozapine. Consistent with previous reports, among these ligands, only roxindole produced any G-protein stimulation from D3R (**Fig. 4B**). Despite this, we found that 3 of these SGA ligands—including roxindole, but also quetiapine and aripiprazole—recruited arrestin-3 (**Fig. 3A, left; 3B-C**). By contrast, clozapine did not produce a change in arrestin-3 localization (**Fig. 3A, right; 3B-C**). These results demonstrate that some SGAs promote arrestin-3 recruitment even in the absence of G-protein signaling (quetiapine, aripiprazole) while others engage neither G-protein nor arrestin-3 (clozapine) at D3R.

**Figure 3:**
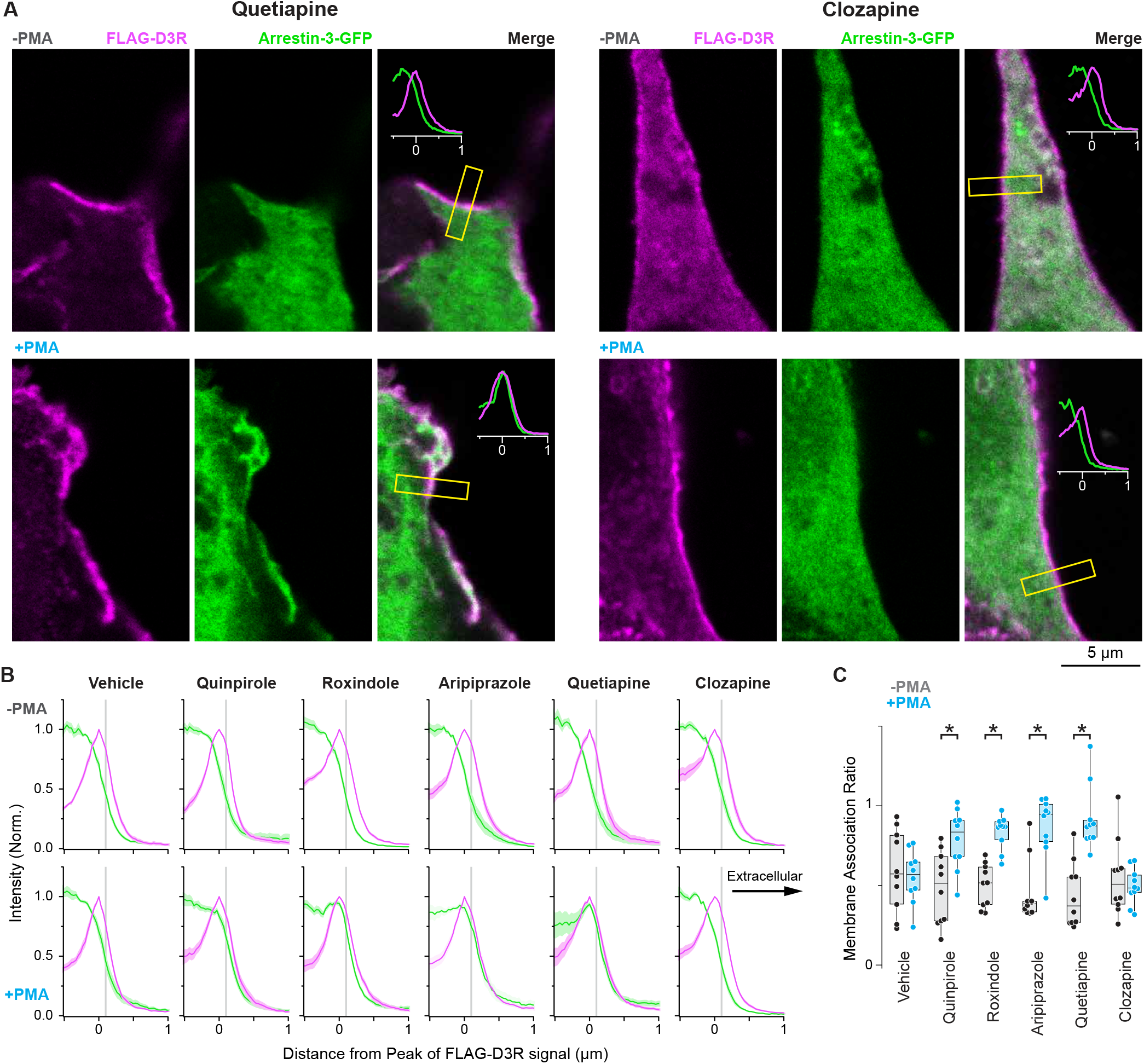
Some SGAs recruit arrestin-3 to D3R. **A**. Example HEK293 cells expressing FLAG-D3R and arrestin-3-GFP. Cells were either treated with drug alone (quetiapine or clozapine), top row, or with drug and PMA, bottom row. Yellow boxes denote regions of analysis. **B**. Quantification of peak arrestin-3 signal relative to D3R signal for each SGA without (top) or with (below) PMA to activate PKC. Each line profile is mean ± SEM. FLAG-D3R in pink, arrestin-3-GFP in green. **C**. Quantification of peak arrestin-3 signal relative to peak D3R signal (membrane association ratio), calculated at the gray bar in B. Black, drug alone; blue, drug plus PMA. Circles represent individual line profiles, 10 lines from 5 cells per condition. *p <0.05, Mann-Whitney test. No change with vehicle or clozapine.

**Figure 4:**
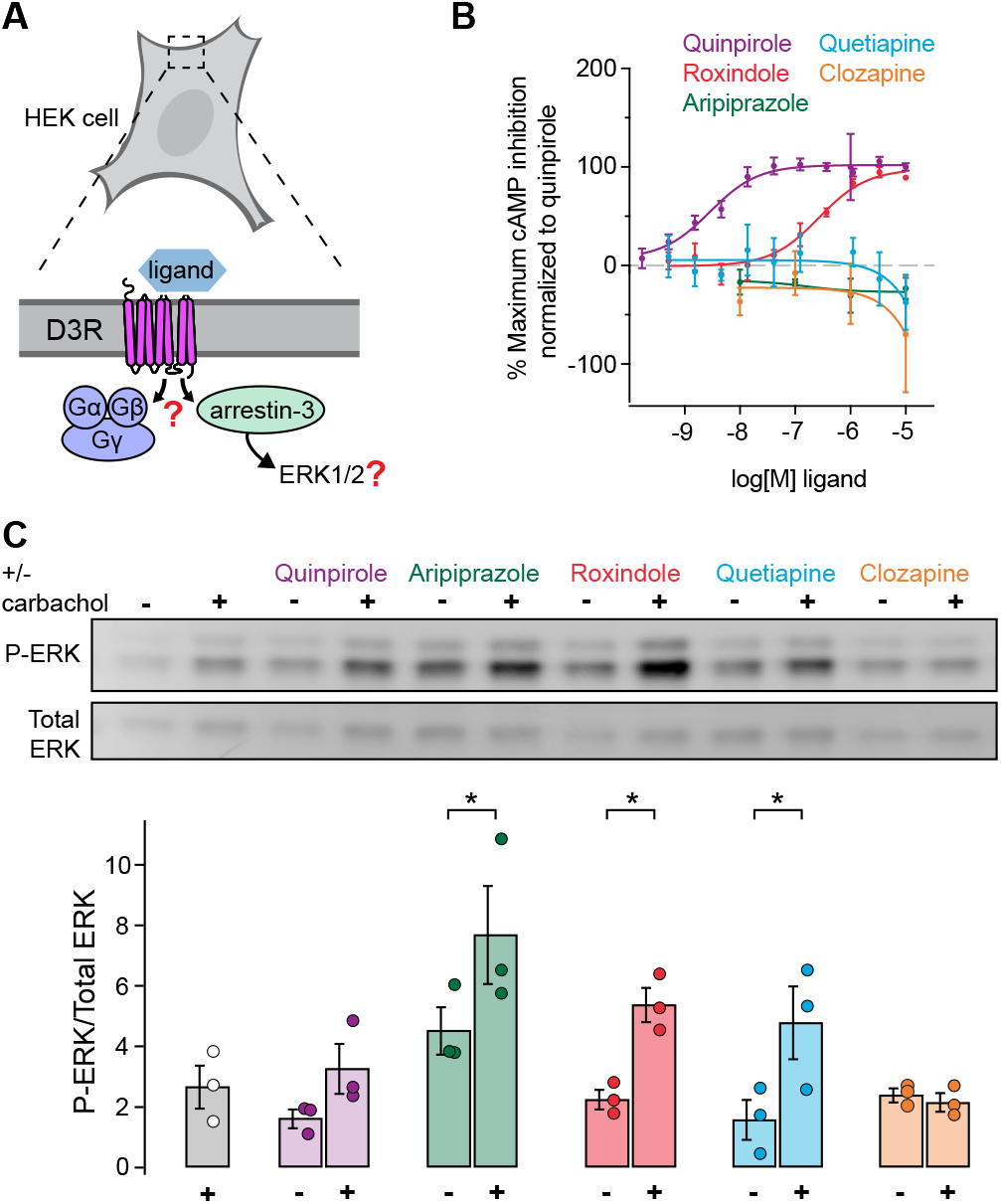
Some SGAs promote ERK phosphorylation through D3R and arrestin-3, not G-protein. **A**. Schematic of experimental question: do SGAs promote ERK phosphorylation? **B**. Percent maximum cAMP inhibition, normalized to quinpirole Emax, as a readout of G-protein signaling for each drug. **C**. Top, representative immunoblot of phospho-ERK and total ERK after 5-minute drug treatment, with or without carbachol to activate PKC, in HEK293 cells stably expressing D3R. Bottom, quantification of phospho-ERK/total ERK across multiple experiments (n=3). Data displayed as mean ± SEM. *p <0.05 compared to drug alone, one-way ANOVA with Holm-Šídák multiple comparisons test.

### Some SGAs promote ERK phosphorylation through D3R and arrestin-3, not G-protein

In a prior study, we showed that both arrestin-3 and ERK are required to modulate Ca_V_3.2 (Yang et al., 2016). ERK phosphorylation by some GPCRs has been shown to occur not only through G-protein activation, but also through arrestin/ERK scaffolding independent of G-protein (Gurevich and Gurevich, 2018). Therefore, we examined the ability of SGAs that did and did not recruit arrestin-3 to promote ERK phosphorylation in HEK293 cells stably expressing D3R (**Fig. 4A**). Given the PKC dependence shown above in Fig. 2 and 3, all experiments were performed in either the absence or presence of PKC activation by carbachol, an agonist for the G_q_-coupled M1 and M3 muscarinic receptors that are expressed endogenously in HEK293 cells. This approach allowed us to titrate a dose of carbachol that stimulated little ERK phosphorylation on its own. Using this approach, we found that the same SGAs that recruit arrestin-3 (**Fig. 3**) also promote ERK phosphorylation but only when PKC is activated (**Fig. 4C**). Specifically, aripiprazole, roxindole, and quetiapine promoted ERK phosphorylation, whereas clozapine did not. These data suggest that some SGAs (e.g., aripiprazole, quetiapine) can promote ERK phosphorylation even in the absence of D3R-mediated G-protein signaling (**Fig. 4B**), classifying them as arrestin-biased agonists. Others (e.g., roxindole) are both G-protein and arrestin-3 agonists, and others still (e.g., clozapine) are antagonists for both.

### Arrestin-biased SGAs modulate AIS calcium in D3+ pyramidal cells in PFC

We established that arrestin-3 is necessary for quinpirole-activated D3R to modulate AIS calcium in PFC (**Fig. 1**). However, quinpirole is both a G-protein and arrestin-3 agonist at D3R, which precluded the ability to determine whether arrestin-3 recruitment alone is sufficient to modulate AIS calcium. The SGAs that promote arrestin-3 recruitment (**Fig. 3**) and ERK phosphorylation (**Fig. 4C**), even in the absence of G-protein activation (**Fig. 4B**), are therefore ideal tools to address whether arrestin-3 recruitment alone can modulate AIS calcium. Whole-cell current-clamp recordings were made from D3+ PFC neurons (**Fig. 1B**), and AP-evoked AIS calcium transients were imaged in the presence of vehicle, quinpirole (positive control), or arrestin-biased SGAs (**Fig. 5A**). These calcium transients were stable for interleaved control cells treated with vehicle for 20 minutes, while 20 minutes of roxindole, aripiprazole, or quetiapine treatment resulted in a ∼30% reduction in AIS calcium compared to baseline, a magnitude comparable to that observed with quinpirole (**Fig. 5B**). By contrast, clozapine did not modulate AIS calcium transients (**Fig. 5B**).

**Figure 5:**
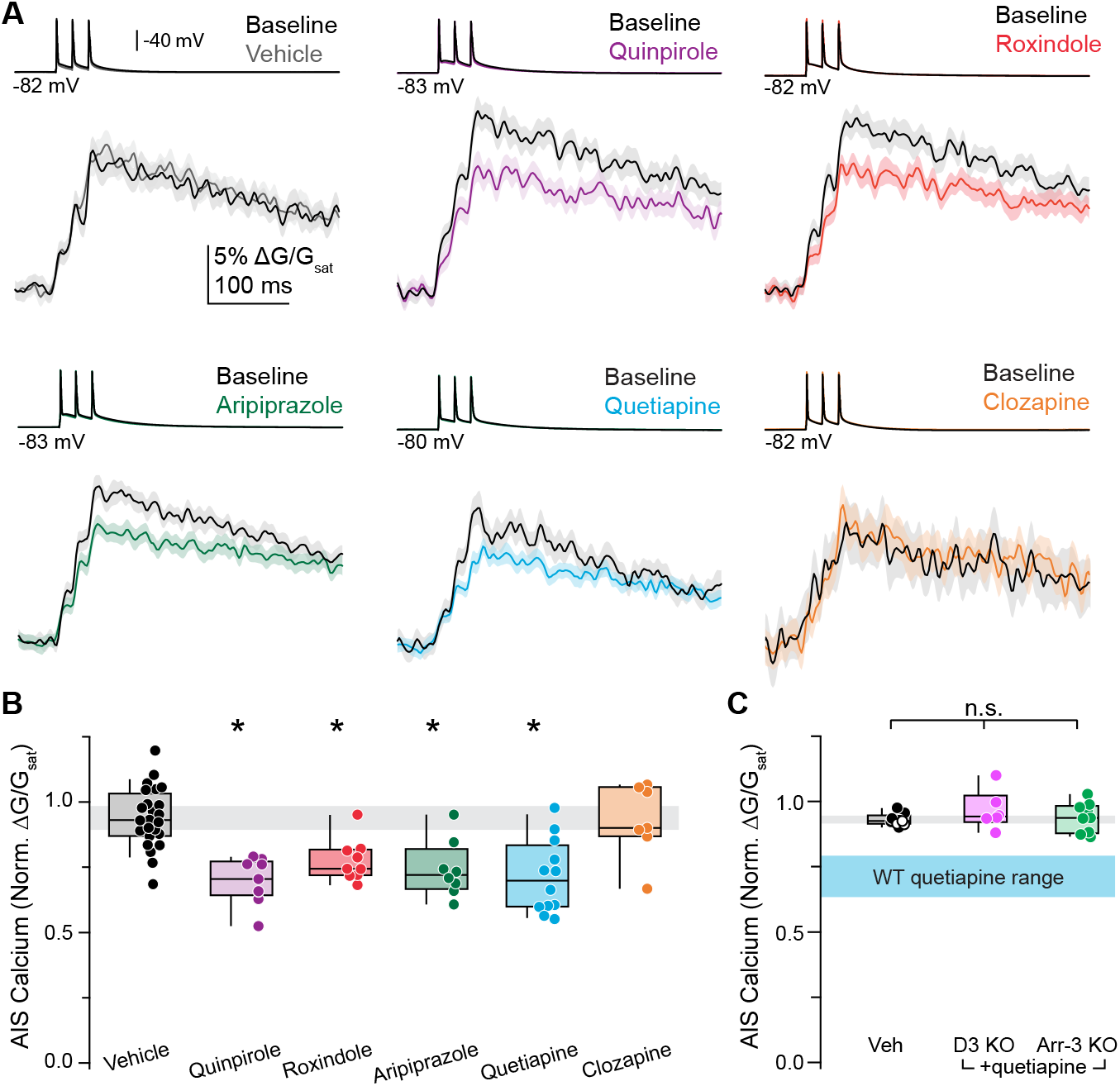
Arrestin-biased SGAs modulate AIS calcium in D3+ pyramidal cells in PFC. **A**. Representative effects of time-locked vehicle control or drug after 20 minutes, averaged across 20 trials per condition. Linescan data displayed as mean ± SEM. Baseline, black; drugs, other colors. **B**. Data summarizing the effects of quinpirole and clinically relevant SGAs on AIS calcium. Gray bar represents 95% confidence interval of control data. Vehicle: 93.1% of baseline, IQR 86.9%-103.3%, n = 24 cells from 21 mice; quinpirole: 70.4% of baseline, IQR 64.2-77.2%, n = 9 cells from 7 mice, p <0.001 (same data as in Fig. 1); roxindole: 74.4% of baseline, IQR 71.9-81.7%, n = 9 cells from 6 mice, p <0.001; aripiprazole: 72.1% of baseline, IQR 66.7-82.0%, n = 8 cells from 6 mice, p = 0.002; quetiapine: 69.9% of baseline, IQR 59.9%-83.5%, n = 12 cells from 6 mice, p <0.001; clozapine: 90.0% of baseline, IQR 86.8%-105.6%, n = 7 cells from 6 mice, p = 0.944. Kruskal-Wallis test with Mann-Whitney U test post hoc (Holm-Šídák correction). **C**. Data summarizing the lack of effect of quetiapine on AIS calcium in D3 KO and arrestin-3 KO mice. Vehicle data include D3 KO (open circles) and arrestin-3 KO (closed circles) mice. Gray bar represents 95% confidence interval of vehicle data from D3KO and arrestin-3 KO animals, while cyan bar represents 95% confidence interval quetiapine data in WT animals. n.s = not significant. Vehicle: 92.5% of baseline, IQR 91.4-94.6%, n = 9 cells from 7 mice; D3 KO: 94.2% of baseline, IQR 92.1-102.3%, n = 6 cells from 3 mice, p = 0.213; arrestin-3 KO: 93.7% of baseline, IQR 87.8-98.3%, n = 9 cells from 4 mice, p = 0.480. Kruskal-Wallis test with Mann-Whitney U test post hoc (Holm-Šídák correction).

SGAs have high affinity for D3R but also bind to other GPCRs. To determine whether the AIS calcium modulation by SGAs occurs via D3R and to demonstrate that SGA modulation of Ca_V_3.2 is arrestin-3 dependent, we assessed AIS calcium modulation by quetiapine in slices from both D3 KO and arrestin-3 KO mice. There was no change in AIS calcium after 20 minutes of quetiapine treatment in D3 KO or arrestin-3 KO animals (**Fig. 5C**). Taken together, these results suggest that some SGAs are arrestin-biased agonists at D3R and can modulate calcium at the AIS in PFC even in the absence of G-protein activation.

### Mice treated chronically with quetiapine, but not clozapine, develop tolerance to the locomotor inhibitory effects of drug

While more prevalent with first-generation antipsychotics, SGAs can also cause locomotor effects, including dystonia and parkinsonism in humans (Casey, 2006; Divac et al., 2014) and locomotor inhibition in mice (Hoffman and Donovan, 1995). While both effects are traditionally thought to be mediated through D2R binding (Divac et al., 2014; Tarsy et al., 2002), mounting evidence suggests that D3R may also be important for locomotion and the locomotor effects of SGAs (Accili et al., 1996; Banasikowski and Beninger, 2012; Botz-Zapp et al., 2021; Bristow et al., 1996; Gyertyán and Sághy, 2004; Kiss et al., 2021; Millan et al., 2004, 1997; Segman et al., 1999; Steen et al., 1997; Svensson, 1994; Waters et al., 1993). Individual SGA have variable degrees of locomotor side effects. We hypothesized that ability to act as an arrestin-3 agonist might contribute to this variability. To examine this hypothesis, we assessed the locomotor inhibitory effects of an SGA that did (quetiapine) and did not (clozapine) recruit arrestin-3. Both quetiapine (15 mg/kg) and clozapine (4 mg/kg) inhibited locomotor activity acutely (**Fig. 6A**), indicating that arrestin-3 recruitment to D3R is not necessary for locomotor inhibition by these SGAs.

**Figure 6:**
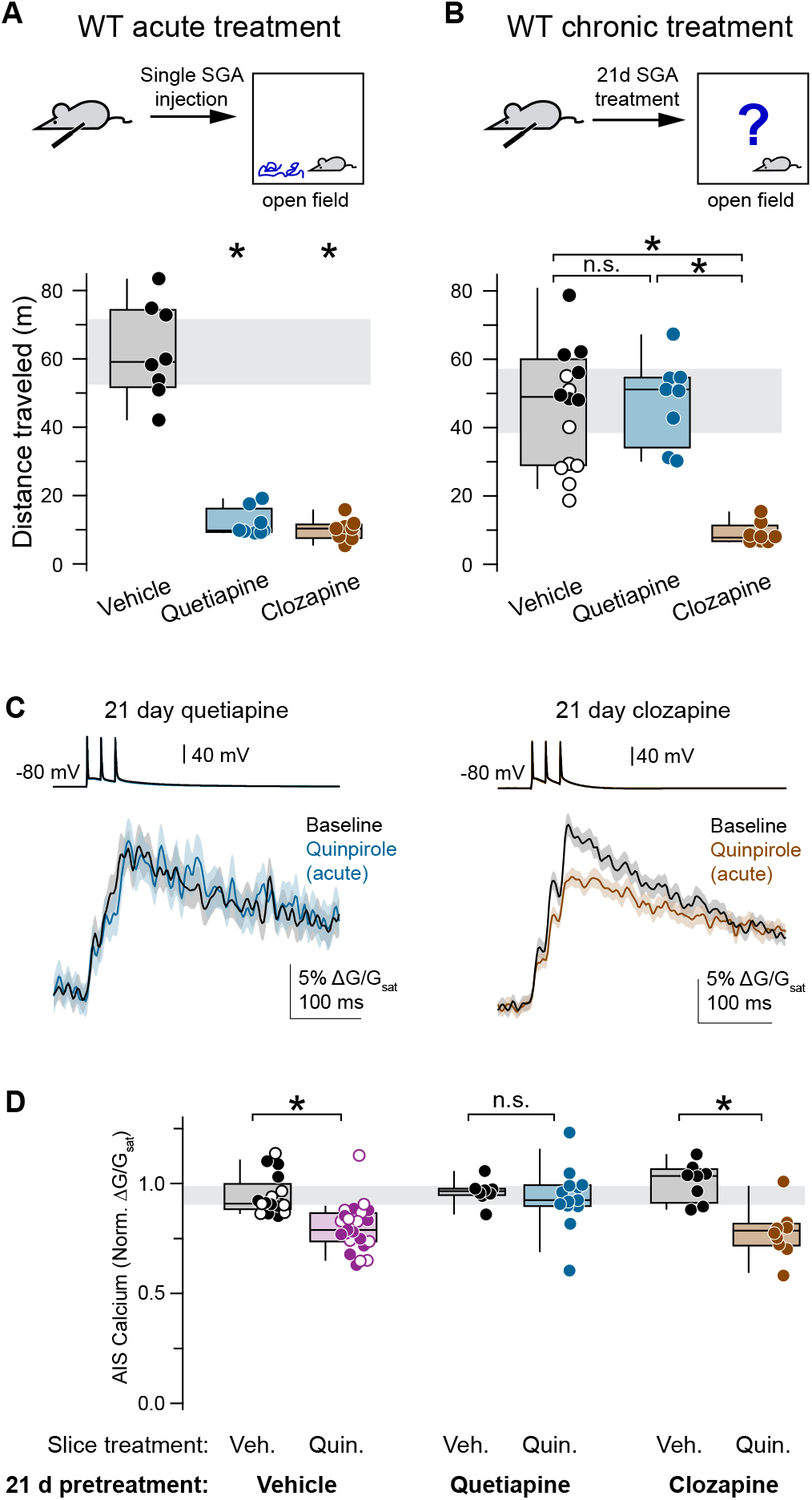
Mice treated chronically with quetiapine, but not clozapine, develop tolerance to locomotor inhibition and are insensitive to D3R-mediated Ca_V_3.2 modulation at the AIS. **A**. Top, mice were injected subcutaneously with drug or vehicle and immediately placed in the locomotor box, and distanced traveled was recorded for 60 minutes. Bottom, total distance traveled for each treatment condition. Gray bar represents 95% confidence interval of vehicle-treated mice. Vehicle: 59.1 m, IQR 51.6-74.3 m, n = 8 mice; quetiapine: 9.81 m, IQR 9.32-16.20 m, n = 8 mice, p <0.001; clozapine: 10.4 m, IQR 7.6-11.6 m, n = 8 mice, p <0.001. **B**. Top, same as A, except mice were pretreated with daily injections of drug or vehicle for 21 days. On day 21, they were placed in the locomotor box immediately following drug injection, and distance traveled was recorded for 60 minutes. Bottom, total distance traveled for each treatment condition. Open circles indicate clozapine vehicle control, while closed circles indicate quetiapine vehicle control. Gray bar represents 95% confidence interval of vehicle-treated mice. Vehicle: 49.0 m, IQR 29.0-60.0 m, n = 16 mice; quetiapine: 51.1 m, IQR 34.1-54.6 m, n = 8 mice, compared to vehicle p = 0.782; clozapine: 7.9 m, IQR 6.8-11.1 m, n = 8 mice, compared to vehicle and quetiapine p <0.001. Kruskal-Wallis test with Mann-Whitney U test post hoc (Holm-Šídák correction). **C**. Representative examples showing effect of quinpirole on AIS calcium transients in mice treated with quetiapine, left, blue, or clozapine, right, brown. Data shown averaged across 20 trials per condition. Linescan data displayed as mean ± SEM. Baseline, black; quinpirole, blue or brown. **D**. Summary data comparing vehicle or quinpirole effect on AIS calcium in animals treated for 21 days with either vehicle, quetiapine, or clozapine. Open circles indicate clozapine vehicle control, while closed circles indicate quetiapine vehicle control. Gray bar represents 95% confidence interval of control data. n.s. = not significant. 21-day vehicle treatment, vehicle slice treatment: 90.9% of baseline, IQR 88.3-99.8%, n = 17 cells from 12 mice; 21-day vehicle treatment, quinpirole slice treatment: 78.8% of baseline, IQR 73.7-86.5%, n = 23 cells from 13 mice, p <0.001; 21-day quetiapine treatment, vehicle slice treatment: 96.5% of baseline, IQR 94.7-97.7%, n = 8 cells from 6 mice; 21-day quetiapine treatment, quinpirole slice treatment: 92.5% of baseline, IQR 89.8-99.3%, n = 13 cells from 6 mice, p = 0.538; 21-day clozapine treatment, vehicle slice treatment: 103.5% of baseline, IQR 91.3-106.6%, n = 8 cells from 4 mice; 21-day clozapine treatment, quinpirole slice treatment: 78.5% of baseline, IQR 71.7-81.7%, n = 10 cells from 6 mice, p = 0.001. Kruskal-Wallis test with Mann-Whitney U test post hoc (Holm-Šídák correction).

While both quetiapine and clozapine inhibited locomotion following acute treatment, the optimal treatment efficacy of SGAs and an equilibrium of effect/side effect profile requires many weeks of treatment, during which additional effects of long-term SGA use can occur. Hence, we also examined the effects quetiapine and clozapine on locomotion following 21 days of drug treatment (15 mg/kg quetiapine or 4 mg/kg clozapine 1x per day for 21 days). Remarkably, on day 21, quetiapine no longer reduced locomotion, while clozapine’s effect was indistinguishable from that observed on day 1 (**Fig. 6B**). These results demonstrate that mice treated chronically with quetiapine develop tolerance to the locomotor inhibitory effects of drug, while animals treated with clozapine do not.

### Chronic treatment with quetiapine, but not clozapine, eliminates D3R-Ca_V_3.2 signaling at the AIS

We next examined whether we could observe drug tolerance on a single cell level. To test this, mice were treated for 21 days with daily injections of quetiapine (15 mg/kg) or clozapine (4 mg/kg) (as in **Fig. 6**). At the end of this regimen, acute slices containing PFC were made, and AIS calcium transients were assessed in D3+ pyramidal cells. In mice treated with quetiapine for 21 days, quinpirole had no effect on AIS Ca_V_3.2 (**Fig. 6C-D**). This lack of quinpirole-induced modulation appeared to occur through quetiapine’s ability to engage arrestin-3, as 21-day treatment with clozapine (or vehicle) did not interfere with quinpirole-mediated modulation of AIS Ca_V_3.2 (**Fig. 6C-D**). These results suggest that chronic treatment with an arrestin-biased SGA, but not an SGA with no arrestin-3 engagement, causes a loss of D3R function at the AIS in PFC. Furthermore, these results, together with the locomotor tolerance phenotype (**Fig. 6A-B**), divide SGA function at D3R into two classes and suggest a possible mechanism underlying the requirement for repeated SGA treatment for maximum benefit.

### Quetiapine-induced loss of D3R-Ca_V_3.2 modulation and development of locomotor tolerance are both mediated by post-endocytic sorting of D3R by GASP1

Recruitment of arrestin-3 to D3R not only arrests the G-protein signal and scaffolds ERK signaling but also promotes receptor endocytosis and subsequent sorting of D3R to the lysosome for degradation through interaction with the GPCR-associated sorting protein-1 (GASP1) (Thompson and Whistler, 2011) (**Fig. 7A**). We therefore hypothesized that repeated treatment with quetiapine would produce tolerance at both the behavioral and single cell level via GASP1 sorting. To examine this hypothesis, we generated mice with a selective disruption of the GASP1 gene only in cells expressing D3R. We accomplished this by crossing D3-Cre driver mice to mice with a conditional floxed GASP1 allele (D3 GASP1 cKO). We then treated these D3 GASP1 cKO mice for 21 days with quetiapine (15 mg/ kg) (as in **Fig. 6, 7**) and measured AIS calcium modulation by quinpirole in D3+ PFC cells. While D3+ neurons from WT mice treated with quetiapine for 21 days showed no quinpirole response (**Fig. 6C-D)**, D3+ neurons from D3 GASP1 cKO mice treated with quetiapine for 21 days showed intact quinpirole-mediated modulation of AIS calcium (**Fig. 7B-C**). To test whether GASP1-mediated post-endocytic sorting of D3R was also necessary for the development of tolerance to the locomotor inhibitory effects of quetiapine, we treated D3 GASP1 cKO mice with quetiapine for 21 days (**Fig. 7D-E**). Unlike WT mice or floxed GASP1 mice not crossed to D3-Cre (21 day vehicle: median 33.9 m, IQR 20.3-45.6 m, n = 4 mice; 21 day quetiapine: median 24.8 m, IQR 16.1-39.6 m, n = 4 mice, p = 0.486, Mann-Whitney test), D3 GASP1 cKO mice did not develop tolerance to the locomotor effects of quetiapine (**Fig. 7D-E**).

**Figure 7:**
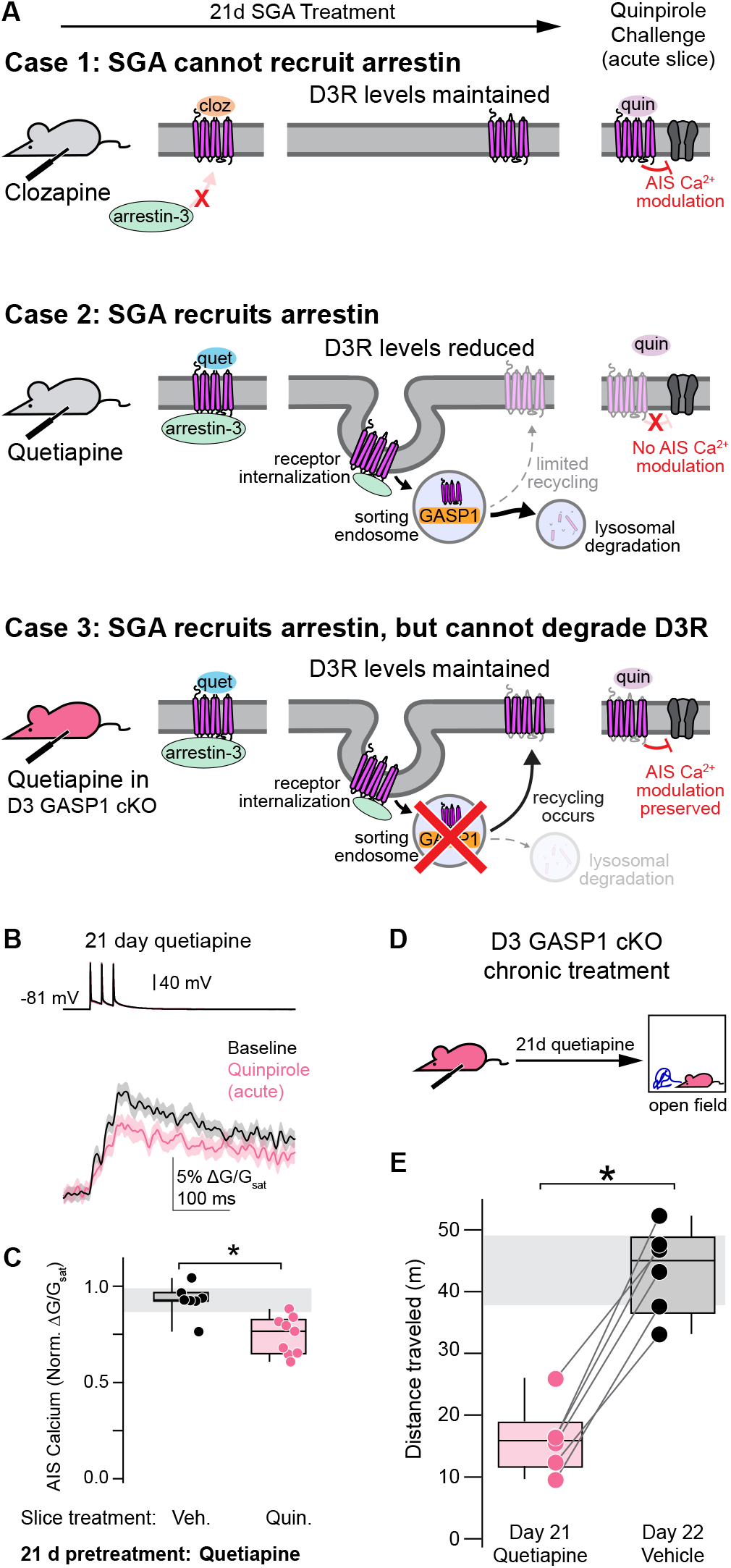
Quetiapine-induced loss of D3R-Ca_V_3.2 modulation and development of locomotor tolerance are both mediated by post-endocytic sorting of D3R by GASP1. **A**. Hypotheses for arrestin-dependent modulation of AIS Ca^2+^ following chronic SGA administration. Clozapine, which does not recruit arrestin-3, does not promote D3R endocytosis, preserving D3R-Ca_V_3.2 modulation (case 1). Quetiapine, which does recruit arrestin-3, promotes D3R endocytosis and degradation, impairing subsequent D3R-Ca_V_3.2 modulation (case 2). Quetiapine administration in GASP1 cKO eliminates lysosomal degradation and is predicted to preserve D3R-Ca_V_3.2 modulation (case 3). **B**. Representative example showing effect of quinpirole on AIS calcium transients in a GASP1 cKO mouse treated with quetiapine. Data shown averaged across 20 trials per condition. Linescan data displayed as mean ± SEM. Baseline, black; quinpirole, blue. **C**. Summary data comparing the vehicle or quinpirole effect on AIS calcium in GASP1 cKO mice treated for 21 days with either vehicle or quetiapine. Gray bar represents 95% confidence interval of control data. 21-day quetiapine treatment, vehicle slice treatment: 92.8% of baseline, IQR 92.3-96.7%, n = 7 cells from 3 mice; 21-day quetiapine treatment, quinpirole slice treatment: 76.6% of baseline, IQR 65.0-82.8%, n = 9 cells from 3 mice, p = 0.005. Mann-Whitney test. **D**. D3 GASP1 cKO mice were treated with quetiapine for 21 days. On day 21, they were placed in the locomotor box immediately following drug injection, and distance traveled was recorded for 60 minutes. The following day, they were injected with vehicle and distance traveled was recorded for 60 minutes. **E**. Total distance traveled for D3 GASP1 cKO mice treated with vehicle or quetiapine. Gray bar represents 95% confidence interval of control data. Vehicle: 45.0 m, IQR 36.5-48.8 m, n = 6 mice; quetiapine: 15.9 m, IQR 11.6-18.8 m, n = 6 mice, p = 0.016, Wilcoxon signed-rank test.

## DISCUSSION

Here, we demonstrate that some SGAs are agonists for arrestin-3 signaling at D3R in the absence of G-protein activity. In D3+ neurons of the PFC, acute slice treatment with these “arrestin-biased” SGAs results in reduced AP-associated AIS calcium influx, demonstrating an acute mechanism of action of a subset of SGAs in addition to their role as D2R and D3R G-protein antagonists. We also establish that chronic treatment with an SGA that recruits arrestin-3 to D3R results in GASP1-mediated post-endocytic sorting to the lysosome and subsequent reduction of both D3R function at the AIS in D3+ PFC pyramidal cells as well as SGA-induced locomotor inhibition. The receptor trafficking-mediated loss of D3R sites of endogenous neuromodulation as a consequence of chronic SGA exposure may be an important aspect of treatment and could help explain why prolonged SGA treatment is necessary for full clinical efficacy.

### Arrestin-3 signaling at D3R

Several studies have focused on arrestin-3 signaling at D2R (Beaulieu et al., 2005, 2007; Kim et al., 2001), and, in the context of antipsychotic drugs, these studies aimed to leverage G-protein or arrestin-3 signaling to maximize therapeutic benefits while minimizing side effects (Allen et al., 2011; Donthamsetti et al., 2020; Masri et al., 2008; Shapiro et al., 2003; Urs et al., 2012, 2016, 2017). Here, we show that select SGAs can also engage arrestin-3 signaling at D3R. We found that in HEK293 cells, some SGAs, with coincident PKC activation, recruit arrestin-3 to D3R. Of note, aripiprazole recruits arrestin-3 to D3R (**Fig. 3**), but not to D2R (Masri et al., 2008), and promotes G-protein signaling at D2R but not D3R. Aripiprazole is thus a G-protein-biased agonist at D2R and an arrestin-biased agonist at D3R. This provides evidence that, even though D2R and D3R are highly homologous, SGAs do not engage the two receptors in the same manner.

Importantly, arrestin-3 recruitment to D3R is sufficient to scaffold ERK phosphorylation even in the absence of G-protein signaling (**Fig. 4**). Specifically, aripiprazole and quetiapine promote increased P-ERK levels (**Fig. 4C**), even though they do not promote G-protein activation from D3R (**Fig. 4B**). This is consistent with a recent study showing that D3R-mediated activation of G-protein and arrestin-3 can phosphorylate ERK on different timescales (Xu et al., 2022). We previously found that in an *ex vivo* slice preparation, quinpirole modulates AIS-localized Ca_V_3.2 channels, hyperpolarizing the voltage-dependence of steady-state inactivation (Yang et al., 2016). This modulation is both arrestin-3- and ERK-dependent (Bender et al., 2010; Clarkson et al., 2017; Yang et al., 2016). We tested here whether “arrestin-biased” SGA agonists at D3R could likewise activate this signaling pathway. We found that only the SGAs that recruit arrestin-3 in HEK293 cells reduce AP-evoked AIS calcium transients in acute slice preparations at timescales consistent with the timing of arrestin-3-dependent ERK phosphorylation [e.g., minutes rather than seconds following ligand application (Xu et al., 2022)].

Kim et al. (Kim et al., 2001) found previously that arrestin-3 did not strongly translocate to D3R in heterologous cells, presumably because there was no concomitant activation of PKC. To our knowledge, D3R is unique in requiring both PKC activation and ligand binding for arrestin-3 recruitment. In HEK293 cells, we achieved PKC activation and ERK phosphorylation by either stimulating PKC directly or by activating a G_q_-coupled GPCR. In neurons, PKC may be activated as a result of ongoing AP activity, as this can induce release of calcium from intracellular stores (Lipkin et al., 2021). In this way, AIS D3R signaling may serve as a coincidence detector for ligand binding and ongoing activity. If this were the case, receptors bound by an arrestin-recruiting dopaminergic ligand would promote signaling only if the neuron was recently firing APs at levels sufficient to activate PKC. Under endogenous conditions, this arrestin recruiting ligand would be dopamine, but dopamine could be substituted with an arrestin-biased SGA during treatment.

Studies of other GPCRs have indicated that arrestin-3 signaling cannot occur in the absence of G-protein activity (Grundmann et al., 2018). For D3R-arrestin-3-Ca_V_3.2 modulation at the AIS, however, our results indicate that D3R can signal through arrestin-3 in a G-protein-independent manner, because SGAs that do not engage G-protein nevertheless produce channel modulation. Even in HEK293 cells, D3R ligands that do not promote G-protein activity recruit arrestin-3 and promote ERK phosphorylation, indicating that this signaling mechanism could be conserved across many cell types. Whether this signaling is unique to D3R remains unknown. While other groups have shown GPCR modulation of ion channels at the AIS (Cotel et al., 2013; Ko et al., 2016; Martinello et al., 2015), D3R’s inhibition of Ca_V_3.2 is, to our knowledge, the first example of a GPCR directly inhibiting a channel solely through an arrestin-3 effector. This finding could have significant implications if it also occurs with other key GPCR drug targets. Future studies should examine ways in which drugs classically considered to be antagonists could likewise promote signal transduction via arrestin-3 to channels or other downstream effectors.

Here, we show that acute application of an “arrestin-biased” SGA to an *ex vivo* slice of mouse PFC modulates Ca_V_3.2 at the AIS via D3R, and that, in a mouse, chronic treatment with the same SGA results in loss of D3R and this modulatory effect. Ca_V_3.2 channels play a critical role in the generation of high-frequency bursts of APs in many cell classes (Molineux et al., 2006). This modulatory pathway decreases the number of Ca_V_3.2 channels that can be recruited during APs, thereby suppressing burst generation (Bender et al., 2010, 2012; Clarkson et al., 2017). Thus, arrestin-3-biased SGAs may acutely act to change bursting properties in D3+ neurons. Upon chronic treatment and loss of membrane D3R, however, neurons may no longer be able to alter their firing patterns based on the presence or absence of dopaminergic input. In PFC, this may be one potential mechanism of SGA action and one that differentiates SGAs into two distinct classes-those that do and do not recruit arrestin-3 and promote loss of D3R function at this site.

Because D3+ pyramidal cells are largely a distinct population of cells from those expressing D1R or D2R in PFC, we were able to isolate SGA-mediated arrestin-3 recruitment at D3R specifically (Clarkson et al., 2017; Gee et al., 2012; Seong and Carter, 2012). The role of D3R signaling in other relevant brain regions, including the hippocampus, nucleus accumbens, Islands of Calleja, and lateral septum, remains unclear (Gurevich and Joyce, 1999; Landwehrmeyer et al., 1993; Prokop et al., 2021; Shin et al., 2018; Suzuki et al., 1998). In future studies, it will be critical to understand fully how D3R regulates neuronal function, either via arrestin-dependent or more canonical signaling pathways, as well as to understand more clearly how these signaling pathways are affected by SGAs (Chen et al., 2006; Diaz et al., 2011; Shin et al., 2018; Swant et al., 2008).

### D3R as a target for SGAs

While modulation of Ca_V_3.2 is an acute SGA effect, antipsychotic drugs often take weeks to months to reach maximal therapeutic benefit. Here, we report two distinct effects of chronic treatment with an arrestin-biased SGA that are mediated by post-endocytic degradation of D3R by GASP1: 1) The ability of dopaminergic drugs, and by extension dopamine, to modulate calcium influx at the AIS is lost, and 2) mice become tolerant to the locomotor inhibitory effects of drug. Specifically, we show that chronic treatment with the arrestin-3-agonist SGA quetiapine causes a profound loss of D3R function at the AIS (**Fig. 6**). This loss is prevented in mice with a disruption of the sorting protein GASP1, which targets GPCRs for degradation instead of allowing their recycling to the plasma membrane, specifically in D3+ neurons (**Fig. 7**). Additionally, we show that chronic treatment with quetiapine results in loss of drug-mediated locomotor suppression (**Fig. 6**), an effect that was also eliminated in D3 GASP1 cKO mice (**Fig. 7**). This result highlights the importance of D3R in the locomotor side-effects of SGAs and, by extension, the effect/side effect profile of SGAs. Together these data directly implicate post-endocytic sorting as a key mechanism controlling the amount of functional D3R in multiple circuits, including cognitive and motor circuitry. Further, these results provide a plausible explanation for why it takes weeks, not hours, for SGAs to show full efficacy. Importantly, this trafficking mechanism efficiently regulates receptor function in a ligand-dependent manner, entirely independent of mRNA expression, potentially reconciling why people living with SMI show dramatically altered receptor protein levels but small, if any, changes in dopamine receptor gene expression (Purves-Tyson et al., 2017).

### Implications for patients

In 2017, roughly 11 million adults, or 4.2% of the adult population, in the United States had a diagnosis of SMI, including bipolar disorder, major depressive disorder, schizophrenia, and schizoaffective disorder (Cohen and Gorrindo, 2020). Recently, several meta-analyses sought to elucidate which antipsychotic drugs most benefit patients (Huhn et al., 2019; McCutcheon et al., 2022). Most people living with SMI rotate through a number of drug treatments and protocols before finding an effective strategy through a protracted and stochastic process. A more complete understanding of the molecular mechanisms underlying the variable acute and chronic actions of these drugs could help inform a more streamlined treatment for an individual patient. Here, we show that SGAs can be separated into two classes based on their bias for D3R-mediated arrestin-3 signaling. SGAs that recruit arrestin-3 can affect D3+ neurons both acutely, by modulating calcium at the AIS, and chronically, by reducing D3R levels at the AIS over time. Historically, GPCR-targeting drugs have been characterized solely for their ability to inhibit or activate G-protein activity. More recently, the role of the arrestin-3-mediated signaling in drug response has come to light. Importantly, arrestin-3 not only arrests G-protein signaling and scaffolds signal transduction to other effectors, including ERK in the case of the D3R, but also promotes receptor endocytosis. Therefore, when considering the implications of arrestin-3 engagement, it is important to account for not only acute signaling but also how engagement can change receptor distribution and surface expression through endocytosis and post-endocytic degradation. This may be particularly relevant for GPCR drugs, such as SGAs, that have rapid pharmacokinetics, reaching the brain within minutes, but that take days or weeks to reach maximal efficacy.

SMI diagnoses describe constellations of symptoms that vary from patient to patient. We posit that for some patients, reducing D3R levels with an arrestin-biased ligand would be most efficacious, while for others, increasing receptor number through blockade of dopamine-mediated arrestin-3 recruitment—and therefore endocytosis and post-endocytic degradation—may be more effective. Moving forward, it would be helpful to track patient diagnosis and symptoms and to map their effect/side-effect profiles onto whether an arrestin-3 biased SGA was or was not therapeutically beneficial. In conclusion, the findings here contribute to a mechanistic understanding of how D3R signaling can vary across different effectors and brain regions and could inform a more personalized approach to treatment with SGAs. D3R has also been suggested as a potential therapeutic target for patients beyond schizophrenia and bipolar disorder, including Parkinson’s disease and substance use disorder (Newman et al.; Van Kampen and Eckman, 2006). Hence, our findings may inform the development of D3R-selective ligands, either biased or not, for multiple indications.

## METHODS

### Experimental model and subject details

All experiments were performed in accordance with guidelines set by the University of California, Davis and University of California, San Francisco Institutional Animal Care and Use Committees. Experiments were performed on wild type or transgenic mice with a C57BL/6 background on a 12-hour light/dark cycle with *ad libitum* access to food and water. Transgenic animals (D3-Cre, D3-Cre::Ai14, D3^-/-^, arrestin-3^-/-^, D3-Cre::Flox GASP1^-/0^) were genotyped by PCR. Transgenic mouse lines had the following research resource identifiers: D3-Cre (KJ302), MMRRC_034696-UCD; D3 KO (D3^-/-^) MGI:4839942; arrestin-3 KO mice were a gift from Robert Lefkowitz and have been backcrossed to C57BL/6 mice for 25 generations. Flox GASP1^tg/0^ mice were created by Cyagen (formerly Xenogen) on a C57BL/6 background. They were generated with the same targeted ES cells as the previously published GASP1-KO mice (see Martini et al., 2010). Briefly, the targeting vector contained 3 lox-P recombination sites, two flanking the GASP1 open reading and a third upstream of a neomycin (G418)-resistance gene inserted into intron 4, which is upstream of the GASP1 open reading frame (ORF) on the mouse X chromosome. A total of 30 μg of NotI-linearized KO vector DNA was electroporated into ∼107 C57BL/6 ES cells and selected with 200 μg/ml G418. Primary ES screening was performed by Southern blotting for the presence of hybridization band corresponding to the targeted allele, and absence of wild-type (WT) hybridization band and a single neo integration. ES clones containing homologous recombination were then transfected with Cre-recombinase and both Type I recombination events, in which only G418 was removed, were identified, and Type II recombination events in which both G418 and GASP1 were identified. Type I events were confirmed on expansion by PCR analysis to create the conditional Flox GASP1^tg/0^ line. Blastocyst injection (using B6/Tyr blastocysts) was performed and the resulting male chimeras were bred with C57BL/6 WT females to generate heterozygous mice.

### *Ex vivo* electrophysiological recordings

Mice aged postnatal day (P) 28-77 were anesthetized, and 250 µm-thick slices containing PFC were prepared. Slices were made from wild-type mice of both sexes in all cases except for experiments using the D3-Cre (KJ302 BAC transgenic) line, as Cre expression is linked to Y chromosome expression in this line.

Cutting solution contained (in mM): 87 NaCl, 25 NaHCO_3_, 25 glucose, 75 sucrose, 2.5 KCl, 1.25 NaH_2_PO_4_, 0.5 CaCl_2_ and 7 MgCl_2_, bubbled with 5% CO_2_/95%O_2_. Following cutting, slices were incubated in the same solution for 30 min at 33°C, then at room temperature until recording. Recording solution contained the following (in mM): 125 NaCl, 2.5 KCl, 2 CaCl_2_, 1 MgCl_2_, 25 NaHCO_3_, 1.25 NaH_2_PO_4_, 25 glucose, bubbled with 5% CO_2_/95% O_2_. Osmolarity of the recording solution was adjusted to ∼310 mOsm. Experiments were conducted at 31-34°C.

Neurons were visualized using Dodt contrast optics for conventional visually guided whole-cell recording or with 2-photon-guided imaging of reporter-driven tdTomato fluorescence overlaid on an image of the slice. All recordings were made from neurons in the upper cortical layers, around the L5a/L5b border, in mPFC. D3R-expressing neurons were identified either using D3-Cre::Ai14 mice and/or were classified as “Type 3” based on intrinsic electrophysiological properties (see Clarkson et al., 2017). Patch electrodes were pulled from Schott 8250 glass (3-4 MΩ tip resistance) and filled with a solution containing the following (in mM): 113 K-gluconate, 9 HEPES, 4.5 MgCl_2_, 14 Tris_2_-phosphocreatine, 4 Na_2_-ATP, and 0.3 tris-GTP at ∼290 mOsm and a pH of 7.2-7.25; 250 µM Fluo-5f and 20 µM Alexa Fluor 594 were added for linescan calcium imaging experiments.

Electrophysiological data were acquired using a Multiclamp 700B amplifier (Molecular Devices) via custom routines in Igor-Pro (Wavemetrics). All recordings were made using a quartz electrode holder (Sutter Instrument) to eliminate electrode drift within the slice. Data were acquired at 20 kHz and filtered at 3 kHz. Pipette capacitance was compensated by 50% of the fast capacitance measured under gigaohm seal conditions in voltage-clamp prior to establishing a whole-cell configuration, and the bridge was balanced. Data were corrected for a 12-mV junction potential. Cells were held around −80 mV, and for linescan experiments, AP were evoked with somatic current injection (3 APs at 50 Hz, 2 nA, 2 ms per stimulus). Time-locked control experiments were interleaved with drug-treated cells daily to ensure stability of recording conditions. Cells were excluded if series resistance changed by >30%, exceeded 20 MΩ, or if Alexa Fluor 594 signal increased over twofold, indicating photodamage of the imaged neurite.

### Two-photon imaging

Two-photon laser-scanning microscopy (2PLSM) was performed as described previously (Bender and Trussell, 2009) using either a Mira 900 with a Verdi 5W pump laser or Chameleon Ultra II laser (Coherent) tuned to 800 nm. Fluorescence was collected with a 40X, 0.8 nA objective paired with a 1.4 NA oil immersion condenser (Olympus). Dichroic mirrors and bandpass filters (575 DCXR, ET525/70 m-2p, ET620/60 m-2p, Chroma) were used to split fluorescence into red and green channels. HA10770-40 photomultiplier tubes (PMTs, Hamamatsu) selected for 50% quantum efficiency and low dark counts captured green fluorescence (Fluo-5F). Red fluorescence (Alexa Fluor 594) was captured using R9110 PMTs.

Fluorescence data were collected in linescan mode, where the laser was repeatedly scanned over a region of the AIS, ∼25-35 µm from the soma, at a rate of 0.5 kHz. The AIS was identified by the absence of spines and its positioning opposite the apical dendrite. Data were averaged over 20 trials and reported as ΔG/G_sat_, which was calculated as Δ(G/R)/(G/R) _max_*100, where G/R_max_ is the maximal fluorescence in saturating calcium (2 mM). Calcium transient peaks were calculated from exponential fits to the fluorescence decay after stimulus offset. Analysis was done in MATLAB, and for presentation, data were smoothed using a 3-point binomial filter in IgorPro.

### Drugs

For HEK293 cell immunostaining, cAMP assays, and immunoblots, aripiprazole and clozapine were purchased from AdooQ Biosciences LLC. For linescan and intraperitoneal (IP) injection experiments, aripiprazole was purchased from Sigma-Aldrich, and clozapine was purchased from Tocris. 5’-*N*-Ethylcarboxamidoadenosine (NECA), quinpirole hydrochloride, and roxindole hydrochloride were purchased from Tocris. Quetiapine hemifumarate was purchased from TCI America. Carbachol and Phorbol-12-myristate-13-acetate (PMA) were purchased from Sigma. Fluo-5F pentapotassium salt and Alexa Fluor 594 hydrazide Na^+^ salt were purchased from Invitrogen. For *in vitro* pGlo and arrestin recruitment studies, drugs were dissolved in DMSO to a concentration of 10 mM and further diluted in water to obtain concentrations between 0.1 nM-10 µM with a final concentration of 0.3% DMSO. For linescan experiments, quinpirole was dissolved in water, while the rest were dissolved in DMSO, to 10 mM (5 mM for aripiprazole) and added to the recording solution for final concentrations of: 2 µM quinpirole, 3 µM roxindole, 2 µM aripiprazole, 1 µM quetiapine, and 1 µM clozapine in 0.01-0.03% DMSO. For vehicle controls, an identical DMSO concentration was added to the recording solution when appropriate. For chronic treatment experiments, quetiapine was given at a dose of 15 mg/kg in 3% DMSO in saline. Clozapine was given at a dose of 4 mg/kg in 4% DMSO in saline. Vehicle controls were 3-4% DMSO in saline, depending on the control condition. For subsequent AIS calcium experiments, injections were given IP; for subsequent open field experiments, injections were given subcutaneously (SC).

### HEK293 Cell Immunostaining

HEK293 cells (ATCC, Manassas, VA) were grown in DMEM with 10% FBS, transfected with D3 (n-terminally tagged with the Flag epitope), arrestin-3-GFP, and GRK 5 using Jetprime transfection reagent (VWR 76299-630) and plated on poly-D-lysine-coated coverslips in a 6-well plate. 24 hours later, live cells were incubated with M1 anti-FLAG antibody (Sigma F3040 1:1000) for 30 minutes at 37°C to label surface receptors. They were then treated with dopaminergic drug (final concentration of 10 µM in 1% DMSO) or vehicle (1% DMSO) for 30 minutes at 37°C and then saline or 10 µM PMA for an additional 30 minutes before fixing. After fixation, cells were blocked and permeabilized in blocking buffer (0.1% Triton-X-100, 3% milk, 1 mM CaCl_2,_ 50 mM Tris–HCl, pH 7.4) and then incubated with fluorescently conjugated secondary antibody (Abcam 150116 1:10000), mounted to glass slides and imaged with a Zeiss LSM 800 confocal microscope (Plan-Apocromat 63x, 1.4 NA) with a pixel size of 0.33 nm using identical exposure and detection settings across all conditions. Cells were tested monthly for mycoplasma.

To quantify relative FLAG-D3R and arrestin-3-GFP localization at the plasma membrane, line profiles (averaged over a 1 µm-wide region) were drawn orthogonal to the membrane, with FLAG and GFP immunostaining plotted relative to each other. Individual fluorescence channels were normalized to the peak of the FLAG-D3R signal and the peak of the GFP signal within 250 nm of the FLAG-D3R peak on the cytoplasmic side. Data are shown as mean ± SEM (shadows) of these line profiles across cells (n = 10 lines per cell, 5 cells per condition). To quantify arrestin-3 recruitment to the membrane, the ratio of GFP to FLAG intensity was calculated at a site 66 nm from the peak of the D3R signal on the extracellular side of the peak.

### Phospho-Erk Assay

HEK293 cells stably expressing FLAG-tagged D3R were grown in DMEM with 10% FBS in 6-well plates coated with 400 µl ECM (Sigma E1270) to 80-90% confluence. Media was then replaced with DMEM with no serum for 16 hours. Media was again removed and replaced with fresh serum-free DMEM for an additional hour. Cells were incubated with drug or vehicle (1% DMSO) for 5 minutes at 37°C (10 µM final concentration of drug ± 1 µM carbachol in 1% DMSO). Cells were lysed in 2x sample buffer containing 1% bromophenol blue, 5% 1M Tris HCl pH 6.8, 1.6% SDS, and 6.4% BME, sonicated for 3 × 10 seconds, and boiled for 5 minutes. Lysates were resolved on a 4-20% acrylamide gradient gel (Thermofisher NP0321BOX) for 3 hours at 100V. Protein was transferred to a PVDF membrane overnight at 4°C with 10% methanol at 30V. The membrane was blocked for 1 hour in 5% BSA, blotted with mouse anti-ERK1/2 (p44/42 MAPK) and rabbit anti-phospho-ERK1/2 (p44/42 MAPK) (Cell Signaling Technologies 4696S and 4370S), both at 1:1000 in 2% BSA at room temperature for 2 hours. The membrane was then washed and immunoblotted with Goat anti-rabbit 800 and Goat anti-mouse 680 (Licor 926-32211 and 926-68070) at 1:10,000 in TBS-T for 1 hour at room temperature in the dark, washed and imaged on the Licor Odyssey Imaging system, and quantified using LI-COR ImageStudio 5 software. Within each blot, data were normalized to the no carbachol, no drug condition.

### D3R-induced cAMP Inhibition

HEK293 cells (ATCC, Manassas, VA) stably expressing both D3R and GloSensor 22F-cAMP (Promega) were created for these experiments. Cells were plated in DMEM + 10% FBS in 384-well white plates at a density of 7,000 cells per well. 24 hours later, media was replaced with CO_2_-independent media + 10% FBS and the GloSensor cAMP reagent (Promega: E1291) at 2% final concentration. Cells were preincubated with drug (0.1 nM-10 µM in quadruplicate) in 1% DMSO for 5 min and then NECA was added (10 µM final concentration) to stimulate cAMP production. The luminescence induced by Glo sensor 22F was measured in real time using a Flexstation3 (Molecular Devices) at 10-minute intervals for 60 minutes at 37°C. The readout at 30 minutes is reported as % NECA-stimulated cAMP activity in the absence of dopaminergic drug and normalized to the quinpirole Emax.

### Open field

Mice were habituated to the locomotion boxes (Med Associates Open Field Arena for Mouse ENV-510 in a sound attenuating box) for 60 minutes the day before their first recorded locomotor session. All recordings were captured and analyzed using Med Associates Activity Monitor 7 Software. Mice were injected SC with drug or vehicle immediately before being placed in the box for a 60-minute recording session. White lights were kept on throughout the recording. Data was recorded in 5-minute bins and is reported as total ambulatory distance (m traveled) per 60 minutes.

### Statistics

All linescan data are reported as medians with interquartile ranges (IQR) in the figure legends and displayed with box plots (medians, quartiles, and 90% tails). All example linescans, as well as intensity plots, are shown as means with standard error shading. Sample sizes were chosen based on standards in the field. No assumptions were made for data distribution, and unless otherwise noted, two-sided, rank-based nonparametric tests were used. Specific statistical tests used are noted in the figure legends. Significance level was set for an alpha level of 0.05, and a Holm-Šídák correction was used for multiple comparisons when appropriate. Statistical analysis was performed using MATLAB, the Real Statistic Resource Pack plugin for Microsoft Excel (Release 8.0), and GraphPad Prism 8 software.

## Acknowledgments

The authors would like to thank Dr. Michael Roberts for help with image analysis, Dr. Matthew McGregor and Anna Lipkin for assistance with data analysis, Anirudh Gaur for guidance on *in vitro* experiments, Dr. Robert Lefkowitz for the arrestin-3 KO mice, Chenyu Wang and Henry Kyoung for assistance with IP injections, and members of the Bender and Whistler labs for critically assessing this work. This work was supported by the National Institute of Mental Health (R01MH112729 to JLW and KJB).

## Author contributions

Conceptualization, S.S., E.L., A.H., K.J.B., J.L.W.; methodology, S.S., E.L., A.H., K.J.B., J.L.W.; formal analysis, S.S., E.L., K.J.B., J.L.W.; investigation; S.S., E.L., A.H., K.J.B., J.L.W.; writing-original draft. S.S.; writing-review and editing, S.S., E.L., A.H., K.J.B., J.L.W.; supervision: K.J.B. and J.L.W.; project administration, K.J.B. and J.L.W.; funding acquisition, K.J.B. and J.L.W.

